# Data-driven gait cycle decomposition based on whole-body coordination dynamics

**DOI:** 10.64898/2026.06.18.732845

**Authors:** Mario De Luca, Matteo Demuru, Enrica Gallo, Marianna Angiolelli, Domenico Tafuri, Giuseppe Sorrentino, Laura Mandolesi, Pierpaolo Sorrentino, Emahnuel Troisi Lopez

## Abstract

The study of human locomotion has long relied on descriptive frameworks of the gait cycle, which have provided essential insights into the functional phases of walking and their underlying biomechanical demands. While these models remain highly informative, they are largely based on observational analyses and may not fully capture the continuous, global coordination that characterizes human movement. The present study proposes an integrated framework to study whole-body coordination. This framework combines network theory with non-negative matrix factorization (NNMF) to treat gait as a dynamic system of coordinated joint interactions. Using three-dimensional kinematic data from 60 healthy subjects, we constructed a representation of whole-body coordination across time, named “dynamic kinectome”. It was then decomposed using NNMF to extract spatial patterns of joint coordination and their corresponding temporal activations, allowing for an interpretable characterization of locomotor organization while preserving physiological meaning. Our analysis extracted six robust, highly consistent, and symmetrical coordination patterns across participants, effectively capturing the primary functional subtasks of locomotion. Rather than challenging classical phase descriptions, these findings enrich them by showing how coordination emerges as a continuous, often proactive process that can extend across conventional phase boundaries and systematically integrates the upper limbs for dynamic stability. Overall, this study provides a holistic, data-driven perspective on human locomotion, offering a promising basis for future investigations in motor control and may contribute to the development of sensitive biomarkers for clinical and rehabilitative applications.

## 1. Introduction

The study of human locomotion is essential across a multitude of disciplines, ranging from clinical rehabilitation to sport science and robotics [1,2]. It provides a fundamental framework for comprehending movement patterns in individuals with and without impairments, thereby offering insight into the intricate neuromotor control mechanisms required for coordination and stability [3]. A gait cycle is traditionally defined as the interval between the initial heel strike and the subsequent ground contact by the same heel. It is evident that this cyclic process is not merely a repetitive mechanical action; rather, it is a highly regulated physiological event, during which several biomechanical phases are carried out. Indeed, as stated by Perry [4], the gait cycle is composed by different phases that are not just a series of timed steps, but a series of problems the body must solve: weight acceptance (i.e. initial contact and loading response), single limb support (i.e. mid-stance and terminal stance), and swing limb advancement (i.e. pre-swing, initial swing, mid-swing and terminal swing). The elegance of Perry’s work lies in the multimodal accuracy with which these phases are described, incorporating data on the biomechanical forces involved (i.e., ground reaction forces and moments), muscular activity (and related synergies), and joint kinematics. However, this subdivision does not take into account the comprehensive whole-body coordination that reflects the harmonious interaction between multiple body segments and muscles, which is essential for producing smooth, stable, and energy-efficient gait patterns [5–7]. Furthermore, without undermining its inherent utility, it is not strictly data-driven as it is mainly descriptive and stems from observational investigations [8]. To overcome the limitations of discrete observational models and in order to study the whole-body coordination, based on previous methodological solutions to similar problems within the electromyography framework [9,10] we adopted a data-driven dimensionality reduction technique consisting of Non-Negative Matrix Factorization (NNMF) [11]. In biomechanics, NNMF is particularly advantageous because its non-negative components are inherently additive, allowing them to be directly interpreted as physiological synergies or motor modules. Initially implemented to extract muscle synergies from electromyographic signals [9,12–14], these mathematical factorization algorithms have been successfully translated to joint angle data to identify continuous kinematic synergies [15–17]. This data-driven decomposition has proven highly effective in demonstrating that the seemingly infinite degrees of freedom of the lower limbs are actually governed by a low-dimensional set of highly reproducible spatiotemporal patterns. However, despite these methodological advancements, the current kinematic factorization literature remains largely constrained by two main limitations. First, analyses are almost exclusively lower-limb-centric, often neglecting the biomechanical role of the upper limbs. Second, traditional approaches typically analyze joint angle trajectories as isolated vectors, failing to capture the multi-segment nature of motor control.

Hence, to obtain a dynamic map of spatiotemporal coordination, we applied network theory to movement data, starting from the kinectome model [18]. In this model, the human body is treated as an interconnected system where nodes represent individual joints and links quantify their kinematic correlation (i.e., coordination). This approach entails a fundamental shift in focus: from analyzing isolated joint trajectories to evaluating the continuous relationships between them [19,20]. To achieve the necessary temporal resolution for gait analysis, we developed a dynamic kinectome matrix capable of capturing these pairwise interactions across the entire gait cycle.

Extracting interpretable patterns from such highly dimensional network data requires robust analytical tools. Hence, we chose a tool capable of reducing the dimensionality of kinematic data and decomposing them in space and time components without compromising the underlying biomechanical meaning.

Therefore, by applying NNMF directly to the dynamic kinectome, this study aims to objectively model the continuous, whole-body multi-joint interactions during walking. This integration of network theory and data-driven factorization facilitates the decomposition of the gait cycle into its fundamental building blocks. Ultimately, this whole-body perspective allows new insights into how the neuromuscular system organizes the continuous flow of human locomotion.

## 2. Methods

### 2.1 Data acquisition and experimental setup

A total of 60 healthy participants (38 males, 22 females; mean age 58.7 ± 12.7) were included in the analysis. Biomechanical assessments were performed at the Motion Analysis Lab (University of Naples Parthenope). To capture kinematic data, an eight-camera stereophotogrammetric system (Pro Reflex Unit; Qualisys Inc., Gothenburg, Sweden) was utilised. Following a modified Davis protocol, 55 reflective passive markers were positioned on the participants’ skin over designated anatomical landmarks [21] (Figure 1). Participants were directed to walk along a linear path at a self-selected, comfortable velocity. For each subject, the dataset comprised two separate recordings. Each recording captured a gait cycle for both the left and right lower limbs, resulting in a total of four gait cycles per participant.

**Figure 1.**
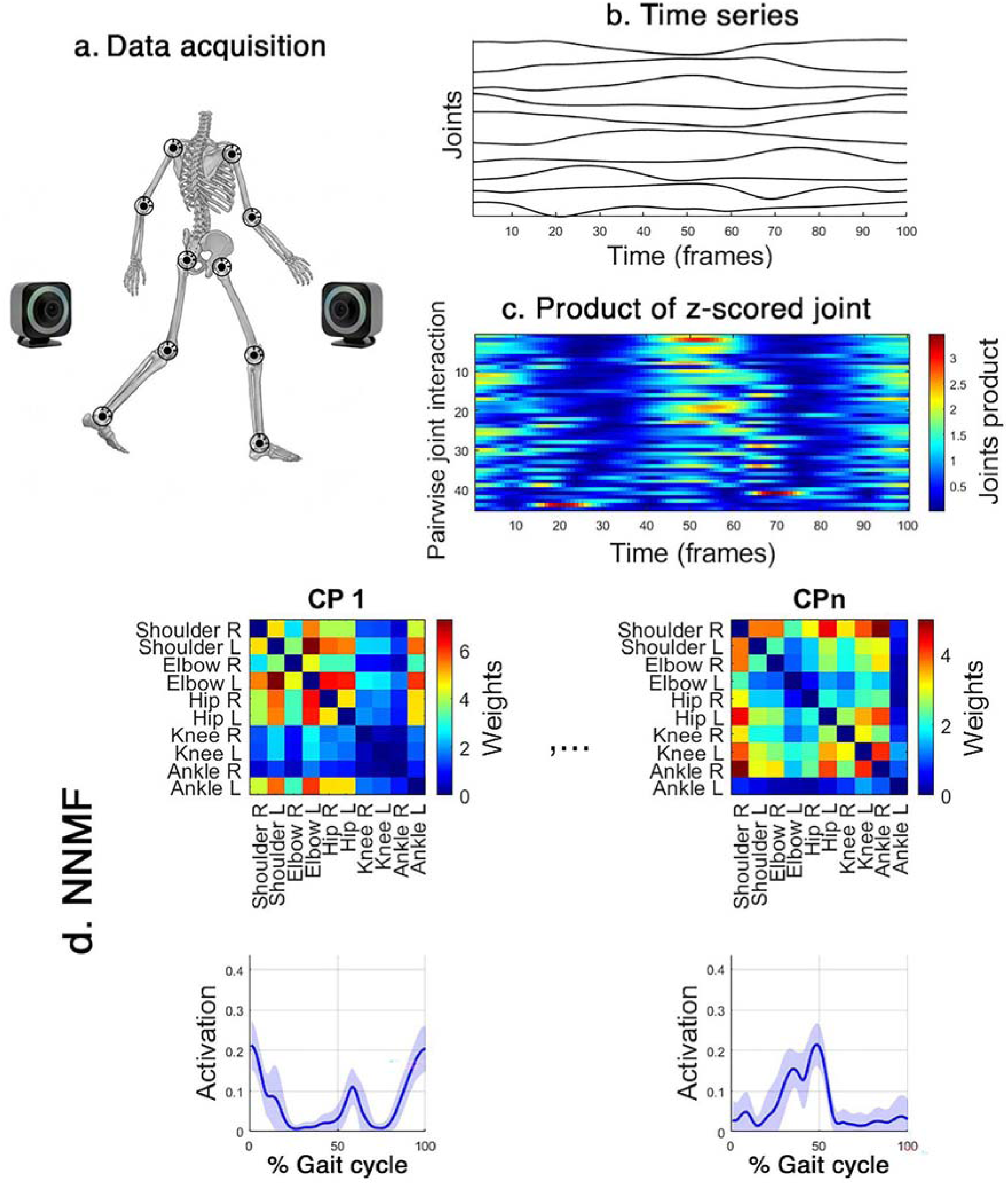
Analysis of the pipeline. (a) 3D kinematics recorded using a stereophotogrammetric system. (b) Data on angular trajectories were extracted and analyzed for ten bilateral joints. (c) Point-by-point product of the z-scored time series for each joint pair to quantify the instantaneous coordination constructing the dynamic kinectome representing pairwise joint interactions over time. (d) The dynamic kinectome matrix was decomposed using the NNMF algorithm to extract spatial coordination patterns and their respective temporal activation across the gait cycle.

### 2.2 Data processing

The Qualisys Track Manager (QTM) software was employed to record and reconstruct the 3D trajectories of the markers. Subsequently, raw data were exported to Visual 3D for preprocessing and the extraction of joint excursion angles. Data was then imported and analysed in MATLAB (MathWorks, version R2024b). As investigating bilateral asymmetries was beyond the scope of this study, all left gait cycles were mirrored to the right side. This approach expanded our dataset to a total of 240 gait cycles. Additionally, to ensure a uniform temporal resolution for the subsequent analyses, all kinematic trajectories were time-normalized via interpolation to 100 data points, corresponding to the length of the shortest recording.

### 2.3 Joint network

To model kinematic coordination, drawing from previous works that have employed network theory to measure motor coordination, we employed a network-based approach to construct a correlation matrix called “kinectome” [18,22]. In this framework, the shoulder, elbow, hip, knee, and ankle joints bilaterally were defined as the network nodes. In the classical kinectome, a single matrix represents a whole time interval enclosing a full movement (e.g., a gait cycle). quantifying coordination via Pearson’s correlation coefficients between the pairs of joints’ time series. To obtain a “dynamic” kinectome over the whole gait cycle, we computed the point-to-point product of the z-scored joint time series. Hence, we obtained a CxT matrix where T is the time axis which contains one hundred time points, and C is the coordination axis that contains the products between all pairs of nodes at each time point.

### 2.4 Non-negative matrix factorization

The NNMF algorithm was used for the dynamic kinectome decomposition into distinct spatial (i.e., coordination pattern) and time-varying components. NNMF is distinguished from the other multi-variate methods by its non-negative constraints [23]. For this reason, we used the absolute values of the dynamics kinectome. Analysis of the explained variance and visual inspection of the components was used to determine the optimal number of components to keep into consideration.

To assess the inter-subject consistency of the extracted coordination patterns, both the spatial and temporal similarities between each individual’s components and the group-averaged templates were quantified using Pearson’s correlation coefficient. Furthermore, to explicitly verify that the extracted spatial synergies represented true physiological coordination rather than mathematical artifacts, the topological results were statistically validated against a spatial null model. For each subject and component, this null model was generated by randomly shuffling the spatial data points, thereby destroying the anatomical network structure while preserving the underlying data distribution. A right-tailed paired t-test was then employed to confirm that the empirical spatial correlations were significantly higher than those derived from the random null model (α = 0.05).

### 2.5 Community detection in coordination patterns

To investigate the modular organization of inter-joint coordination (the “kinectome”), community detection was performed using the Louvain algorithm optimized through a consensus modularity approach[24,25]. Rather than computing communities on the grand-average connectivity matrix, which risks obscuring individual topological features, the analysis was carried out on each participant’s connectivity matrix independently for each of the 6 movement patterns (10 × 10 kinectomes × 60 subjects per pattern). For each subject and pattern, the community assignment vector (membership vector) was extracted. To quantify the topological consistency of these community structures across the cohort (N = 60), we computed the Normalized Mutual Information (NMI) for all pairwise subject combinations, yielding a 60 × 60 similarity matrix for each movement pattern. The global consistency of a pattern was defined as the mean off-diagonal NMI value.To assess whether the observed group-level consistency was statistically significant or emerged by chance, a permutation test (10000 permutations) was implemented. For each permutation, node assignments were shuffled independently for each subject, and the surrogate mean NMI was recomputed to build an empirical null distribution and extract *p*-values. Finally, to extract a single, representative group-level partition for each movement pattern without losing individual topological information, an association (co-occurrence) matrix was constructed. This 10 × 10 matrix stores the probability (from 0 to 1) that any pair of joints belongs to the same community across the 60 subjects. Elements with a co-occurrence probability lower than 0.30 were thresholded to minimize background noise. The final representative group-level communities were extracted by applying the consensus Louvain algorithm to this group-association matrix [24].

### 2.6 Statistic

To evaluate the robustness of the identified gait patterns across the lifespan of our cohort, we employed Pearson’s correlation coefficients to examine the relationship between chronological age and the spatial and temporal similarities of individual gait cycles relative to the group mean. Specifically, we correlated the sum of similarity values for spatial and time components, separately.

Furthermore, to determine if age-related physiological changes necessitated a partitioned analysis, we performed a sensitivity analysis by stratifying the cohort into age-specific subgroups. The stability of the coordination topology was assessed across these strata to ensure that the global network metrics were not confounded by age. Since the correlation strength was moderate and no significant topological breakthroughs were observed between groups, all subsequent network analyses were performed on the unstratified, pooled sample to maximize statistical power.

## 3. Results

### 3.1 Dimensionality reduction

The dynamic kinectome matrices derived from the joint coordination were decomposed using non-negative matrix factorization algorithms. The optimal number of components was determined by analysing the variance explained as a function of the number of extracted components and the visual inspection of the results. While mathematical heuristics such as the elbow method on the explained variance suggested a lower-dimensional subspace (N = 4), visual inspection and biomechanical considerations of the gait cycle indicated that this number merged functionally distinct gait phases. Previous literature highlights that purely variance-based criteria may under-factorize physiological datasets, masking fine motor control mechanisms. Therefore, we fixed the number of components to N = 6, including components until their time intervals are not overlapped. Notably, the temporal profile of the first component exhibited equivalent activation at both the beginning (0%) and the end (100%) of the gait cycle. This dual presence reflects the periodic nature of the movement, as the defined time window initiates and terminates with the exact same kinematic event (i.e., initial contact). Furthermore, statistical validation against a null model confirmed that all six extracted components were highly consistent across the 240 gait cycles (p < 0.001), with a correspondence between individual and average pattern equal to 85% for spatial patterns and 83% for temporal patterns.

### 3.2 Temporal patterns

As shown in the figure 2, the six extracted components display interesting temporal correspondences to the well known phases of the gait cycle, representing the “when” of joint coordination. Particularly, the first component (0-10% of the cycle) shows peak activity during the *initial contact* and *loading response* and again during the terminal swing (86-100%), suggesting a role in weight acceptance and preparation for ground contact. The second component (11-21%), overlaps half of the *mid stance phase*. The third component (22-36%) goes from the second half of the mid *stance phase* to the beginning of the *terminal stance phase*. The fourth component (36-60%) mainly corresponds to the *terminal stance phase* and *pre-swing phase*. Then, the fifth (61-72%) and the sixth (73-85%) components mostly overlap with the initial swing, and the mid swing phases, respectively.

**Figure 2.**
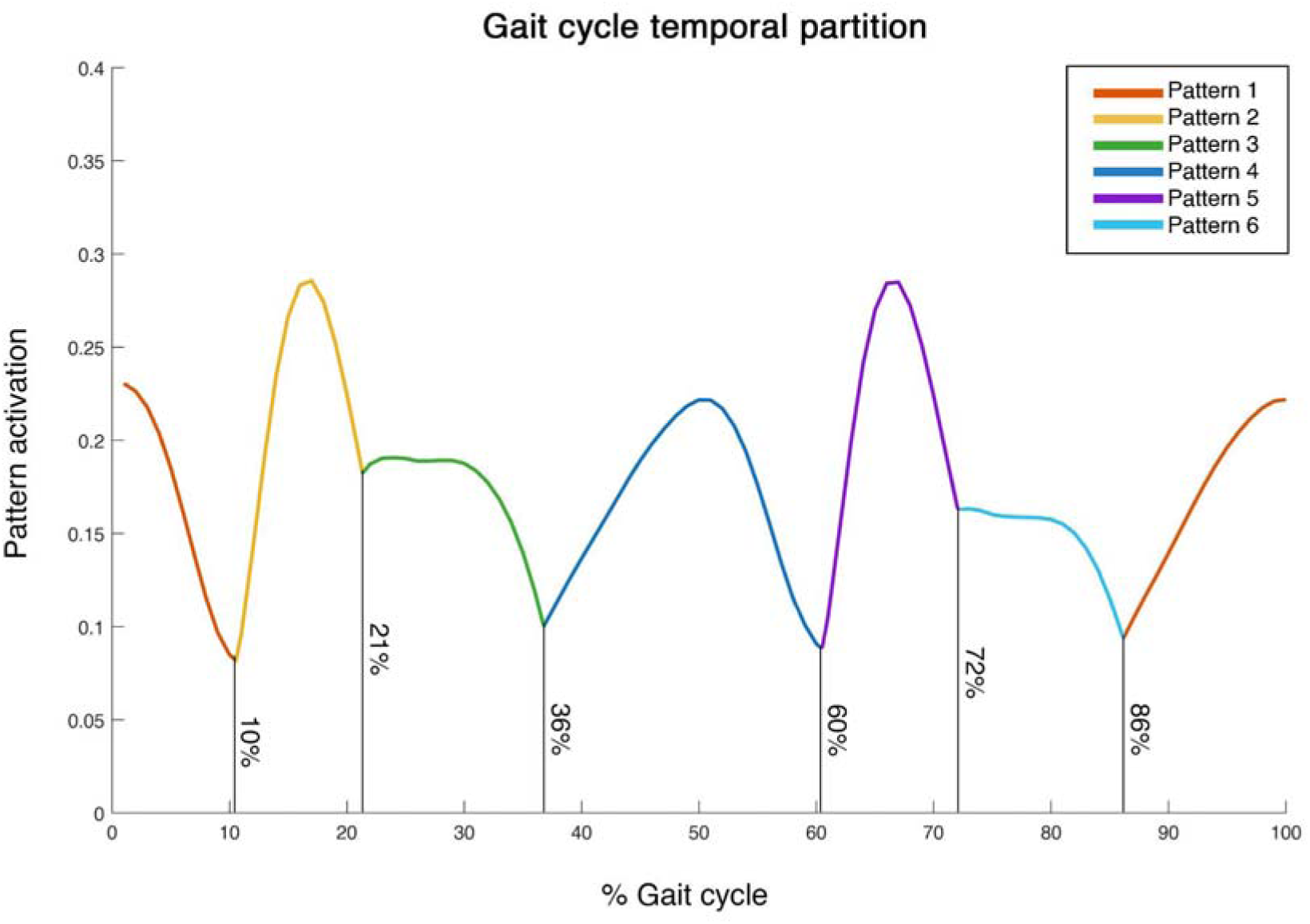
Gait cycle temporal partition. The plot displays the temporal activation profiles of the components extracted via non-negative matrix factorization (NNMF). The continuous curves are graphically segmented at their intersection points, highlighting the specific interval during which each pattern exhibits dominant activation. Consistent with an alternating gait, the analysis revealed two sets of three symmetric patterns. Due to the continuous cyclical nature of locomotion, the activation profile of the first component wraps around the 0% and 100% boundaries, which correspond to the initial foot strike.

**Figure 3.**
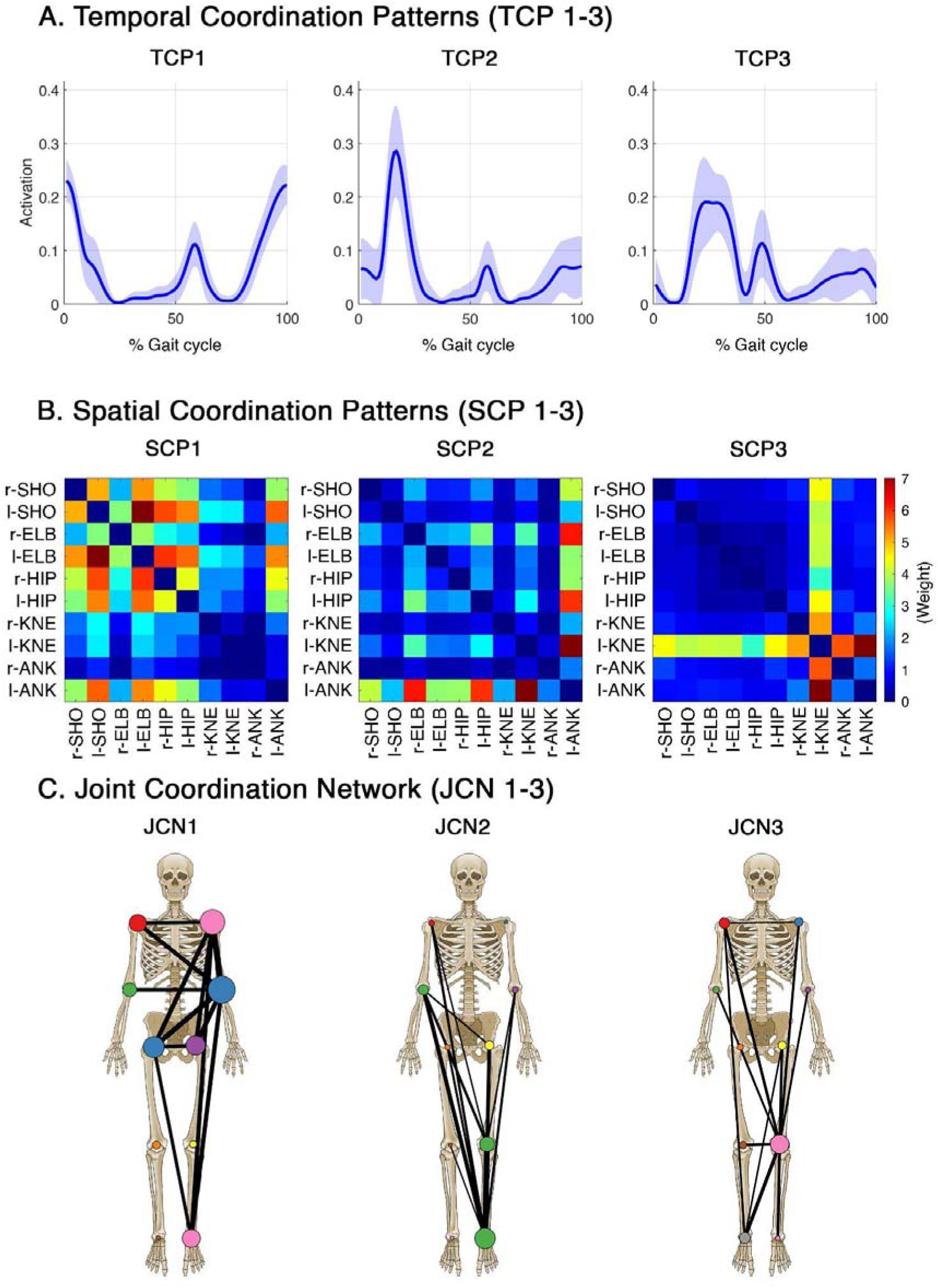
Spatiotemporal coordination patterns and network representations (pattern 1-3). (A) Temporal Coordination Patterns (TCP 1-3) illustrating the mean activation profiles (solid lines) and standard deviations (shaded areas) across the normalized gait cycle. (B) Spatial Coordination Patterns (SCP 1-3) depicted as kinectome heatmaps, representing the coupling weights between pairs of joints. (C) Joint Coordination Networks (JCN 1-3) mapped onto an anatomical skeletal model for direct visual interpretation. For visual clarity, only the top 25% strongest connections (edges) are plotted. Nodes are colored according to the consensus Louvain clustering algorithm to highlight distinct functional joint communities. l-: left; r-: right; SHO: shoulder; ELB: elbow; KNE: knee; ANK: ankle.

### 3.3 Spatial patterns

The spatial patterns define the “how” of the decomposition, illustrating the specific joint couplings that dominate each phase. These matrices represent the coordination between ten joints bilaterally: shoulder, elbow, hip, knee, and ankle. These patterns reveal six distinct and symmetric coordination patterns, which were highly consistent across participants when averaged at the group level (p < 0.001). Although the recordings were segmented to capture a complete gait cycle of the reference limb, the contralateral limb simultaneously executes its own concurrent cycle. This bilateral nature of walking is naturally captured by the NNMF decomposition, which identified six distinct coordination patterns that emerge in pairs, corresponding to the two sides of the body.

Figure 3 presents the first three patterns extracted via NNMF:

- The first coordination pattern (NMI = 0.858, p < 0.001) emerges at the onset and conclusion of the right gait cycle (i.e., during the loading response and terminal swing phases). It is characterized by two synergies: the first one includes the left shoulder, and the left ankle, while the second one includes the left elbow, and right hip.

Furthermore, this pattern exhibits a broader contribution from the upper limb joints (both shoulders and elbows), the hips, and the left ankle, which is preparing for foot-off.

- The second coordination pattern, occurring at the beginning of the right mid-stance phase, is driven by a single dominant synergy (MNI = 0.939, p < 0.001). In this instance, the coordination network primarily comprises the left ankle, left knee, and right elbow, while the functional contribution of the remaining joints is minimal.

- The third coordination pattern, which occurs during the second half of the right mid-stance, shows a single synergy (MNI = 0.919, p < 0.001), as well, that includes the left knee and the left ankle. However, beyond the sinergies, the main contribution to this pattern is given by the left knee, which accomplishes its extension task through a highly distributed coordination strategy, coupling almost equally with all other joints.

Consistent with the alternating and symmetric nature of human locomotion, the remaining three components (patterns 4, 5, and 6), presented in Figure 4, emerged as perfectly specular replications of the first three patterns. Specifically, these components exhibited an identical spatiotemporal structure, but with a complete left-right inversion of the involved joint communities and a temporal phase shift corresponding to the contralateral step:

**Figure 4.**
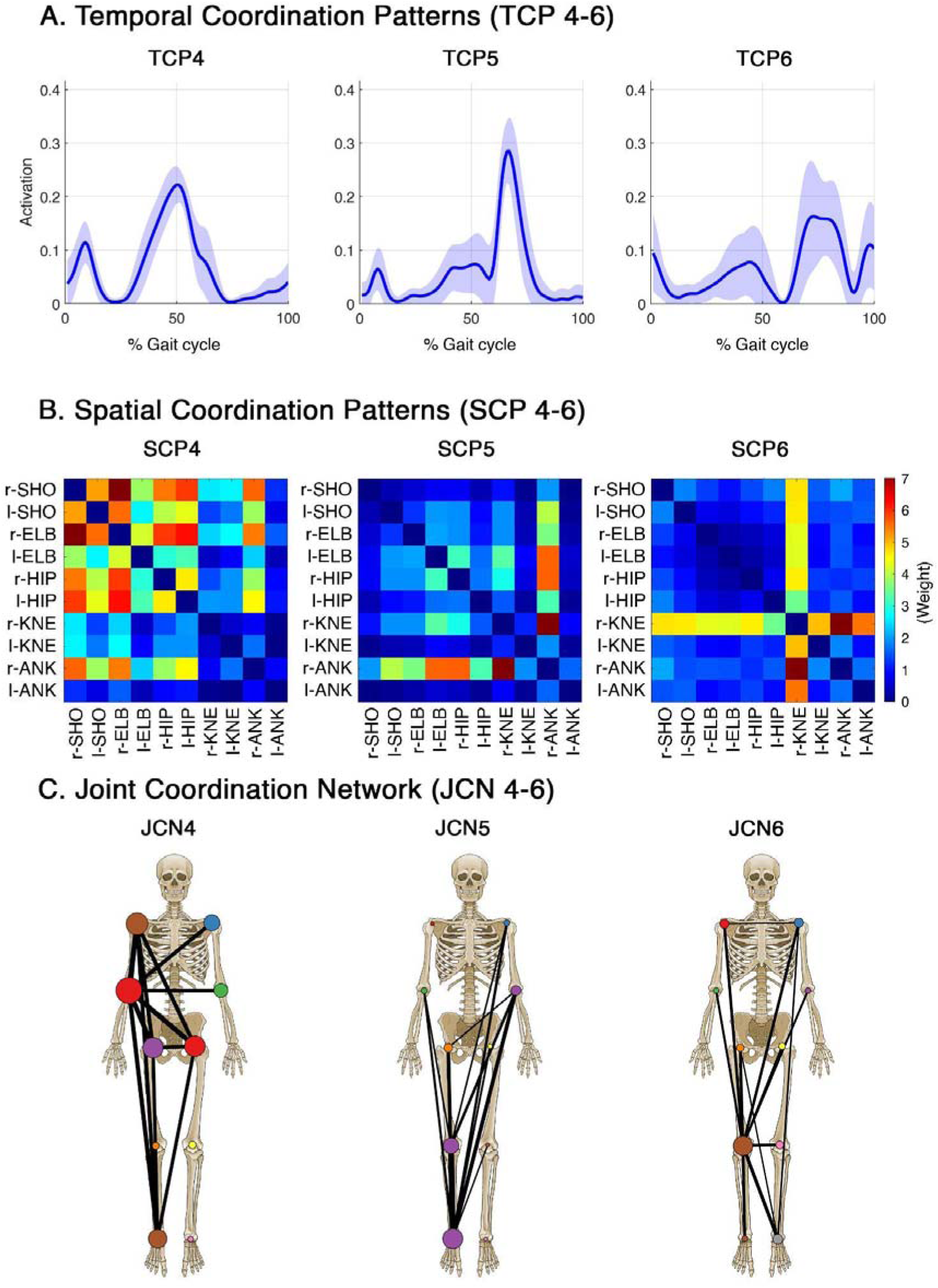
Spatiotemporal coordination patterns and network representations (pattern 4-6). (A) Temporal Coordination Patterns (TCP 4-6) illustrating the mean activation profiles (solid lines) and standard deviations (shaded areas) across the normalized gait cycle. (B) Spatial Coordination Patterns (SCP 4-6) depicted as kinectome heatmaps, representing the coupling weights between pairs of joints. (C) Joint Coordination Networks (JCN 4-6) mapped onto an anatomical skeletal model for direct visual interpretation. For visual clarity, only the top 25% strongest connections (edges) are plotted. Nodes are colored according to the consensus Louvain clustering algorithm to highlight distinct functional joint communities. l-: left; r-: right; SHO: shoulder; ELB: elbow; KNE: knee; ANK: ankle.

- Pattern 4 (MNI = 0.864, p < 0.001): synergy 1 includes right shoulder and right ankle; synergy 2 includes left elbow and right hip.

- Pattern 5 (MNI = 0.945, p < 0.001): synergy 1 includes right elbow, left knee, and left ankle.

- Pattern 6 (MNI = 0.927, p < 0.001): synergy 1 includes right knee and right ankle.

Furthermore, an overall assessment of the temporal activation profiles revealed two prominent kinematic features. First, the topological involvement of the upper body joints was systematically concentrated during, or immediately adjacent to, the double support phases. Second, the temporal boundaries of the extracted patterns did not strictly align with classical gait events; notably, the foot strike event did not mark the onset of a new coordination pattern, but rather occurred midway through an ongoing kinematic synergy that consistently initiated during the preceding terminal swing phase.

Furthermore, we found that the age of the participants was inversely correlated to both spatial (r = -0.30; p = 0.019) and temporal (r = -0.31 p = 0.015) similarity of the subjects to the group-averaged coordination patterns. That means that the spatiotemporal structure of coordination proportionally changes while aging. Nevertheless, since these age-induced variations were minimal and stratifying the cohort by age groups yielded no major topological or temporal shifts, we decided to proceed with the unstratified whole sample. The core spatiotemporal structure of the gait cycle remains sufficiently conserved across ages, ensuring that our downstream network analyses are not negatively affected.

## 4. Discussion

In this study, we sought to build upon the established observational decomposition of the gait cycle proposed by Perry, by applying dimensionality reduction techniques, specifically NNMF, to objectively decompose a large kinematic dataset into its fundamental constituent parts [4]. By applying this algorithm to the dynamic kinectome, we successfully leveraged whole-body joint coordination to achieve a purely data-driven decomposition [26]. This approach elevates the analysis from isolated joint kinematics to an interconnected network model, yielding a comprehensive, informative, and objective representation of the human gait cycle. Ultimately, our analysis produced a highly consistent 6-component model that effectively captures the primary functional subtasks of locomotion.

When comparing current knowledge in gait cycle studies with our findings, one of the main results concerns the temporal misalignment between the extracted kinematic synergies and classical clinical gait events [27]. Traditionally, the foot strike is considered the starting point of the gait cycle and the onset of the weight acceptance phase. However, our results showed that, from a whole-body coordination perspective, this event does not mark the beginning of a new motor pattern. Instead, the foot strike occurs midway through the first coordination pattern, which consistently initiates during the preceding terminal swing. This temporal shift suggests that the central nervous system employs a proactive, anticipatory kinematic strategy rather than a purely reactive one. By initiating the coordination network prior to ground contact, the body pre-configures its joint kinematics in anticipation of the need to absorb the impact and manage the transition into weight acceptance [28].

Equally noteworthy is the topological composition of these coordination patterns, particularly regarding the substantial involvement of the upper limbs. Clinical gait analysis is commonly, but not exclusively, lower-limb-centric. Our kinectome analysis, which incorporated not only the upper limbs but also the whole-body spatiotemporal relationships, revealed that the upper joints (i.e., shoulders and elbows) are systematically coupled with the lower body during, or immediately adjacent to, the double support phases [29,30]. The double support phase represents a critical period of dynamic instability [31], characterized by the transfer of the body’s center of mass from the trailing limb to the leading limb. The active recruitment of the upper body network during these specific temporal windows highlights the role of arm-swing and inter-limb coordination in counteracting angular momentum, fine-tuning dynamic balance, and ensuring a smooth, energy-efficient transfer of weight [32]. Bridging the initial weight acceptance and the single-limb support, the second coordination pattern (and its specular fifth counterpart) highlights a highly specific diagonal synergy.

Occurring at the onset of the stance phase for the reference limb, this network is primarily driven by the contralateral lower limb (i.e., knee and ankle) and the ipsilateral elbow. From a biomechanical point of view, while the reference limb begins to bear the body’s weight, this specific spatiotemporal coupling suggests the combination of two tasks. The coordinated flexion of the swinging knee and ankle reflects the foot clearance, which is simultaneously counterbalanced by the opposite arm, to regulate the angular momentum and allow the forward progression [33].

As the reference limb progresses into the second half of mid-stance, the third coordination pattern (and its specular sixth counterpart) captures a fascinating kinematic shift. While the stance limb provides vertical support, the contralateral knee plays the role of a kinematic hub as it extends forward during its swing phase. Interestingly, the kinectome revealed that this swinging knee does not operate within a specific cluster of more joints; rather, it exhibits a highly distributed coordination strategy, coupling almost equally with all other joints. From a biomechanical point of view, the forward extension of the swinging limb generates inertial forces and angular momentum. To prevent these forces from destabilizing the body’s center of mass over the single supporting foot, the entire kinematic chain must dynamically counterbalance the swinging leg. This ubiquitous coupling suggests that managing the inertial perturbation of the swing phase requires continuous, whole-body postural integration, anchored by the trajectory of the advancing knee [34].

Following the completion of these first three fundamental subtasks, the coordination sequence intrinsically repeats for the contralateral side. As extracted by the NNMF, patterns four, five, and six emerge as almost perfectly specular replications of the first three. This high degree of spatiotemporal symmetry is particularly noteworthy because the factorization algorithm was entirely unsupervised and uninformed by anatomical lateralization. Therefore, this mirrored recurrence serves as a robust proof of concept for both the methodological reliability of the kinectome and the physiological integrity of the analyzed healthy cohort [18]. From a neurophysiological perspective, establishing this highly symmetric, six-component baseline, may provide a standardized reference framework to potentially identify and quantify asymmetric coordination deficits in future clinical populations.

Finally, our analysis revealed a significant inverse correlation between participant age and their spatiotemporal adherence to the group-averaged coordination patterns. From a physiological perspective, this gradual divergence aligns with the known neuromotor adaptations associated with healthy aging [35]. Older adults often exhibit subtle alterations in gait kinematics, such as modified joint excursions, cautious stabilization strategies, or altered sensorimotor integration [7,36–38]. Consequently, these alterations may translate into deviations from the ideal spatiotemporal motor pattern. However, the fact that these variations are continuous and do not result in a complete topological shift underscores the robustness of our 6-component model. The core spatiotemporal framework of locomotion is highly conserved across the adult lifespan. Aging induces gradual, fine-grained modulations within these synergies, rather than a fundamental reorganization of the central motor plan, confirming the generalizability of the kinectome approach across different age demographics.

In summary, the spatiotemporal factorization of the dynamic kinectome characterizes the organizational framework of human coordination, reflecting the operational principles of the neuromuscular system. By representing locomotion as a continuous sequence of six distributed coordination patterns, this framework may recall the higher-order motor planning employed by the central nervous system to simplify the control of complex, multi-joint tasks. A noteworthy outcome of this decomposition is the relative absence of tightly clustered patterns dominating the weight-bearing limb during its stance phase. We hypothesize that this phenomenon inherently stems from the strictly kinematic nature of our input data. The stance limb, constrained by ground reaction forces, is primarily governed by isometric stabilization and kinetic demands rather than large joint excursions. Consequently, the kinematic variations are naturally skewed towards the unconstrained swinging limb and the upper body, which actively manage angular momentum and postural balance. Rather than a limitation, this characteristic highlights the unique strength of the kinematic network approach. By organically filtering out the rigid, force-driven mechanics of the constrained limb, our methodology successfully isolates and reveals the coordination motor strategies. Ultimately, the kinectome paradigm offers a robust, objective, and holistic tool to decode human locomotion, paving the way for advanced applications in motor control research and clinical rehabilitation.

Several limitations of the present study should be acknowledged. First, to establish a foundational baseline without unnecessary analytical complexity, all left-sided gait cycles were mirrored to the right, precluding the analysis of limb dominance or bilateral asymmetries, which were not within the scope of the paper. Second, while age-related correlations emerged, our sample size prevented a robust age-stratified analysis. Future studies with larger cohorts should investigate these age-induced adaptations, while strictly controlling for variations in walking speed. Finally, as our current network model relies exclusively on joint kinematics, future research should transition toward a multimodal approach, integrating electromyographic and kinetic data, to fully map the underlying neuromuscular drive onto its spatial mechanical execution.

In conclusion, our data-driven and coordinative 6-component model partially mirrors the functional subdivision traditionally described in gait analysis. Through these differences, our approach integrates previous knowledge on the topic and demonstrates that gait phases are not merely descriptive time bins, but are driven by highly specific and time-varying joint networks. By treating the body as a dynamic network, computing the equal kinectome, and combining it with the NNMF, it is possible to capture the continuous nature of human locomotion. This framework offers significant highlights also in the clinical field. Since these components represent a baseline of healthy and efficient neuromotor control, any deviations in either the spatial networks or temporal activations could serve as sensitive biomarkers for pathological gait. Future studies should apply this pipeline to populations with neurological or orthopedic impairments to identify exactly where, and how the coordinative network breaks down.

## Acknowledgements

This research was funded by European Union “NextGenerationEU”, (Investimento 3.1.M4. C2), EBRAINS-Italy of PNRR, grant number IR0000011 and Governo Italiano Ministero per lo sviluppo Economico, ACCORDI PER INNOVAZIONE. Approccio User-friendly integrato per Diagnosi, Assistenza e Cura Efficaci—AUDACE grant number B69J23006050007.

## Author contribution

E.T.L., M.D.L. conceptualized the study; M.D.L., E.T.L., E.G., M.D. performed experiments; E.T.L., M.D.,M.D.L., analyzed the data; G.S., D.T., L.M., P.S. supervised the study; M.D.L., M.D., E.T.L. wrote the manuscript; G.S.,P.S., E.T.L, L.M. reviewed and edited the manuscript.

## Data Availability Statement

data are available on reasonable request to the corresponding author.

## Institutional Review Board Statement

the study was conducted in accordance with the Declaration of Helsinki, and approved by the Ethics Committee of Psychological Research of the Department of Humanities of the University of Naples Federico II (protocol code 26/2020 approved on 10/9/2020).

## Informed Consent Statement

Informed consent was obtained from all subjects involved in the study.

